# Novel MAFA+, ISL1+, NKX6-1-stem cell derived pancreatic cell clusters secrete insulin and control blood glucose in rodent models of diabetes

**DOI:** 10.1101/2023.10.20.563345

**Authors:** J. Ratiu, S. Southard, W. Rust

**Affiliations:** Seraxis, Inc

## Abstract

This article describes a stem cell line derived by reprogramming of native human islet cells that consistently generates pure populations of endocrine pancreatic clusters following a simple differentiation protocol. Surprisingly, the population of stem cell derived pancreatic endocrine clusters that was most consistently capable of regulating blood glucose in rodent models of diabetes lacked robust expression of the key beta cell maturation-associated factor NKX6-1 but did manifest high expression of other key drivers of endocrine cell specification and maturation, ISL1 and MAFA. These data support the hypothesis that multiple pancreatic profiles can be identified in stem cell derived cultures and that these have disparate in vivo potency. The population with low NKX6-1 and high in vivo potency was further characterized by transcriptome profiling as an endocrine-committed population progressively maturing in vitro to a state proximal to the native islet.

## Introduction

The islet replacement cure for type 1 diabetes is unavailable to most patients due to lack of donor islet supply and the requirement for potent immune suppression [1]. Islet-like clusters grown from immortal human pluripotent stem cells are a viable replacement for cadaveric islets and can be manufactured at large scale [2]. Pluripotent stem cells are defined by an inherent ability to follow maturation pathways that specify every tissue of the developing embryo and, eventually, all adult tissues. Protocols for guiding pancreatic maturation of pluripotent stem cells aim to trigger pancreatic fate choice and restrict non-pancreatic lineages. These protocols create populations comprised of cells with functional pancreatic features and off-target cells [3-5]. We hypothesize that the presence of cells without appropriate endocrine specification can diminish the clinical potency and safety of the population.

Induced pluripotent stem cells are epigenetically distinct from embryonic stem cells and can retain epigenetic imprinting of the source tissue, leading to differential gene expression and differentiation capacity between cell lines [6]. We aimed to capitalize on this phenomenon by reprogramming primary pancreatic islet cells from a consented human donor to create a stem cell line with enhanced pancreatic differentiation potential. To achieve this, we developed a protocol for episomal reprogramming of these primary pancreatic islet cells and then searched among the immortalized clones for populations that most efficiently responded to maturation cues to re-form populations of cells with pancreatic hallmarks [7]. The best performing cell line (clone SR1423) is characterized by a restricted fate potential and therefore failed to meet criteria of pluripotency. Further characterization reported here emphasizes the distinction between SR1423, induced pluripotent stem cells (iPSCs), and embryonic stem cells (ESCs).

Stem cell-derived pancreatic clusters are less functionally mature than primary islets and contain cell populations with transcriptomic and epigenomic profiles that share some features with fetal, neonatal and infant cell populations [8]. To date, the fates of differentiating cell populations that express subsets of pancreatic markers are not well defined. In particular, distinct progenitor populations are identified based on differential expression of the transcription factor NKX6-1 among cells expressing the transcription factor PDX-1 in vitro and in vivo [8, 9]. PDX-1 is a key determinant of pancreatic identity that is activated early in pancreatic commitment and remains expressed through development and in the mature endocrine and exocrine pancreatic tissue [10]. NKX6-1 is activated in early pancreatic endocrine cell commitment and is again expressed in insulin-secreting beta cells [11]. Among populations of differentiating pluripotent stem cells, it has been suggested that PDX-1+/NKX6-1+ are fated to form functional insulin-secreting beta cells, while PDX-1+/NKX6-1-cells form polyhormonal cells that either resolve in vivo to glucagon-secreting alpha cells or non-functional off-target cells [3, 11-14]. NKX6-1 also has an important role in the specification of the enterochromaffin population of serotonin-producing enteroendocrine cells of the gut [15]. Pancreatic and enterochromaffin populations expressing NKX6-1 emerge from a common NGN3-expressing progenitor population and share similar functional features [3, 5, 15, 16]. Because serotonin-expressing cells are common among populations of stem cell-derived pancreatic cells, it is speculated that stem cell-derived pancreatic populations contain off target cells with enterochromaffin identity [16-19]. More recent analyses of transcriptome profiles of the developing human fetus reveal that pre-beta cells also express serotonin and that the serotonin-expressing population among differentiating stem cells that was identified as enterochromaffin may be bona fide progenitors of pancreatic islet cells [8]. The expression of transcription factors ISL1 and MAFA are also expressed in early endocrine progenitors and again in maturing islets, indicating roles in both specification of endocrine-committed progenitors and in maturation or maintenance of functional endocrine cells [9]. It is unclear which set of transcription factors reflect the highest in vivo potency for islet replacement therapy.

We designed two maturation protocols that both guided the differentiation of SR1423 cells to clusters that contained native proportions of C-peptide and glucagon-expressing cells, and demonstrated robust, physiologic glucose responsive C-peptide secretion. C-peptide, a by-product of insulin synthesis is secreted by beta cells in equimolar concentrations to insulin and is frequently used as a reliable surrogate marker of insulin production and secretion [20, 21]. The population defined by the first protocol was characterized by a high proportion of cells expressing the transcription factors PDX-1 and NKX6-1, and C-peptide. The second protocol primarily generated cells that lacked expression of NKX6-1 but maintained expression of PDX-1 and C-peptide. Second protocol cells more robustly expressed the pancreatic transcription factors ISL-1 and MAFA than cells from the first protocol.

Clusters of cells derived from each protocol were implanted to rodent models of type 1 diabetes. Both types of clusters created stable grafts in the hosts with persistent expression of PDX-1, chromograninA and other hallmarks of pancreatic endocrine cells. Over time, however, the NKX6-1-high population (first protocol) lost expression of C-peptide and failed to consistently regulate blood glucose. Grafts of the NKX6-1-low clusters (second protocol) maintained C-peptide expression, consistently regulated blood glucose and activated expression of NKX6-1 in vivo. We selected the second protocol NKX6-1-low clusters that demonstrated high in vivo potency and developed cGMP manufacturing processes to support IND-enabling nonclinical studies and clinical studies. In-depth transcriptome analysis of the in-process manufactured cell populations revealed a progressive maturation leading directly towards the transcriptome of the native adult islet.

## Results

### Undifferentiated stem cell line

The human cell line, SR1423, was generated by causing transient expression of the Yamanaka factors in cells of human islets harvested from a consented donor pancreas with high HLA and blood type compatibility for allogeneic donor organ transplantation [7]. Rather than screen for pluripotency, this cell line was selected for its ability to differentiate toward the definitive endoderm and, subsequently, PDX-1-expressing pancreatic progenitor cell fate. Analysis showed that SR1423 does not meet criteria for pluripotency as it fails to express the master control gene for mesoderm specification, brachyury, in response to mesoderm-inducing agents [7]. The hPSC Scorecard assay was employed for a comparison of pluripotency-related genes to a reference dataset of multiple pluripotent stem cells [22]. Undifferentiated SR1423 demonstrated a gene expression profile outside of the range of the reference dataset with mesodermal differentiation potential scoring lowest (Figure S1). Consistent with previous directed differentiation experiments, the T gene (TBXT, brachyury) was highly down-regulated while the early endoderm specifying transcription factors SOX17 and HNF1B was found to be up-regulated without any external differentiation cues [7].

Transcriptome analysis by single-cell RNA sequencing (scRNA-seq) also demonstrated a gene expression profile of the undifferentiated cells that manifests distinctions from previously published iPSC and ESC datasets (figure 1) [23, 24]. Unsupervised clustering with iPSC (GSE202398) and hESC (GSE143783) single cell datasets revealed SR1423U are unique from each stem cell type but more similar to iPSC than hESC (Fig 1A). Pathway analysis also demonstrated that transcriptional regulation of SR1423U falls between that of iPSC and hESC (Fig 1B) with a unique profile of transcription factor expression (Fig 1C). Consistent with hPSC Scorecard Assay data and lack of mesoderm potential, low expression of POU5F1 (OCT4), NANOG and LIN28 suggest incomplete reprogramming of SR1423U cells [25]. Together, these data demonstrate that SR1423U is a unique, partially reprogrammed islet-derived stem cell.

**Figure 1.**
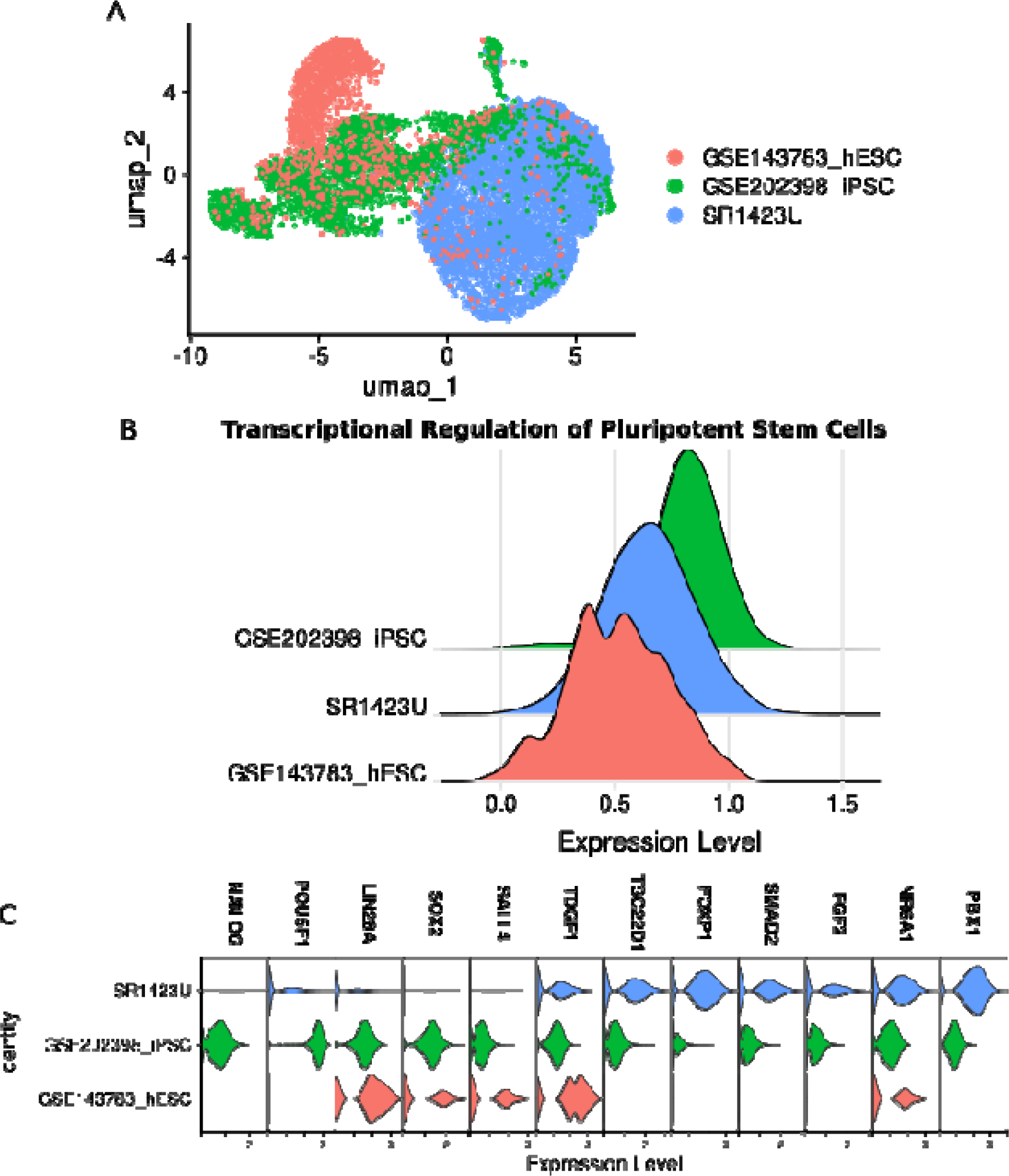
SR1423 are unique stem cells. (A) UMAP embedding of SR1423U, H1 hESC, and iPSC cells analyzed by snRNA-seq. (B) Ridge plot of the gene signature for transcriptional regulation of pluripotent stem cells. (C) Violin plot showing expression of key stem cell transcription factors in SR1423, hESC, and iPSC cells.

### SR1423 differentiation

SR1423 responds to basic differentiation protocols that drive endocrine pancreatic fate choice and generates highly pure populations of cells with pancreatic endocrine cell characteristics [7]. As previously described, SR1423 is a multipotent, and not pluripotent stem cell line. We therefore characterize SR1423 as a multipotent stem cell line with a pancreatic endoderm bias. SR1423 was derived and maintained in a cleanroom following good manufacturing practice (GMP) principles. Master cell and working cell banks were created and characterized for lack of adventitious agents and other parameters relevant for clinical application. Most work reported here was performed on aliquots from the working cell bank.

First differentiation protocol: A review of the literature that describes the differentiation of pancreatic islet cells from pluripotent stem cells suggest that an ideal target population contains a large proportion of cells that co-express the transcription factors PDX-1 and NKX6-1, and the hormone insulin, a profile found in the beta cells of the native adult islet [4, 18, 26]. We therefore designed a modified version of our previously published protocol based on these principles that drives SR1423 to generate islet-like clusters with a high proportion of cells that expressed PDX-1, NKX6-1, and the by-product of insulin processing, C-peptide [7] (detailed in methods, table 1). Flow cytometry and immunofluorescent staining (figure 2) showed a population of cells that co-express PDX-1, NKX6-1 and C-peptide. These cell populations secreted C-peptide in response to glucose in vitro (figure 3). C-peptide expression is found in the absence of NKX6-1, and some NKX6-1 expressing cells lack C-peptide (figure 2). Thus, NKX6-1 expression is neither necessary nor sufficient for C-peptide expression in this cell population.

**Table 1.**
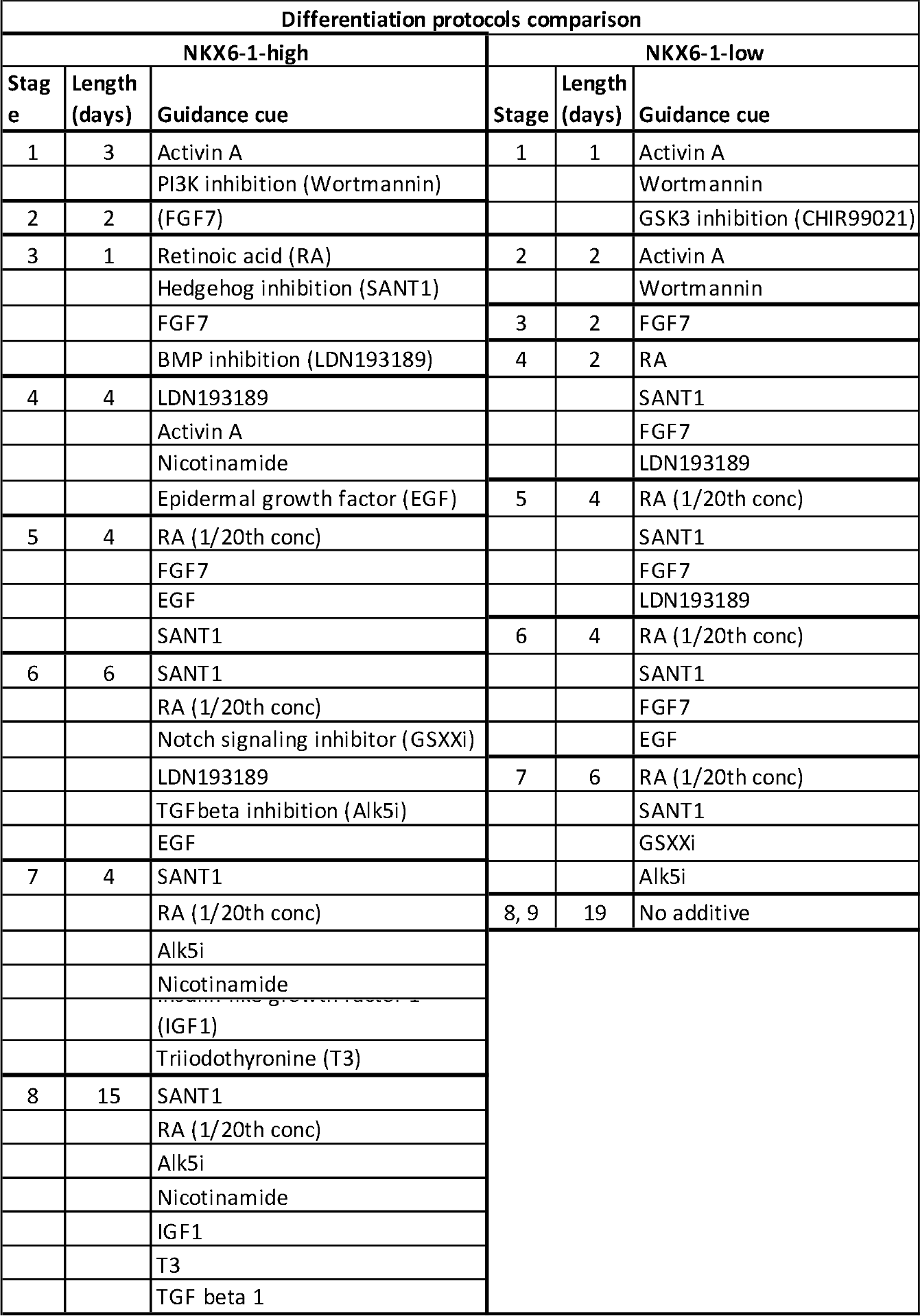
Comparison of guidance cues that drive differentiation of SR1423U to NKX6-1 high and low populations.

**Figure 2.**
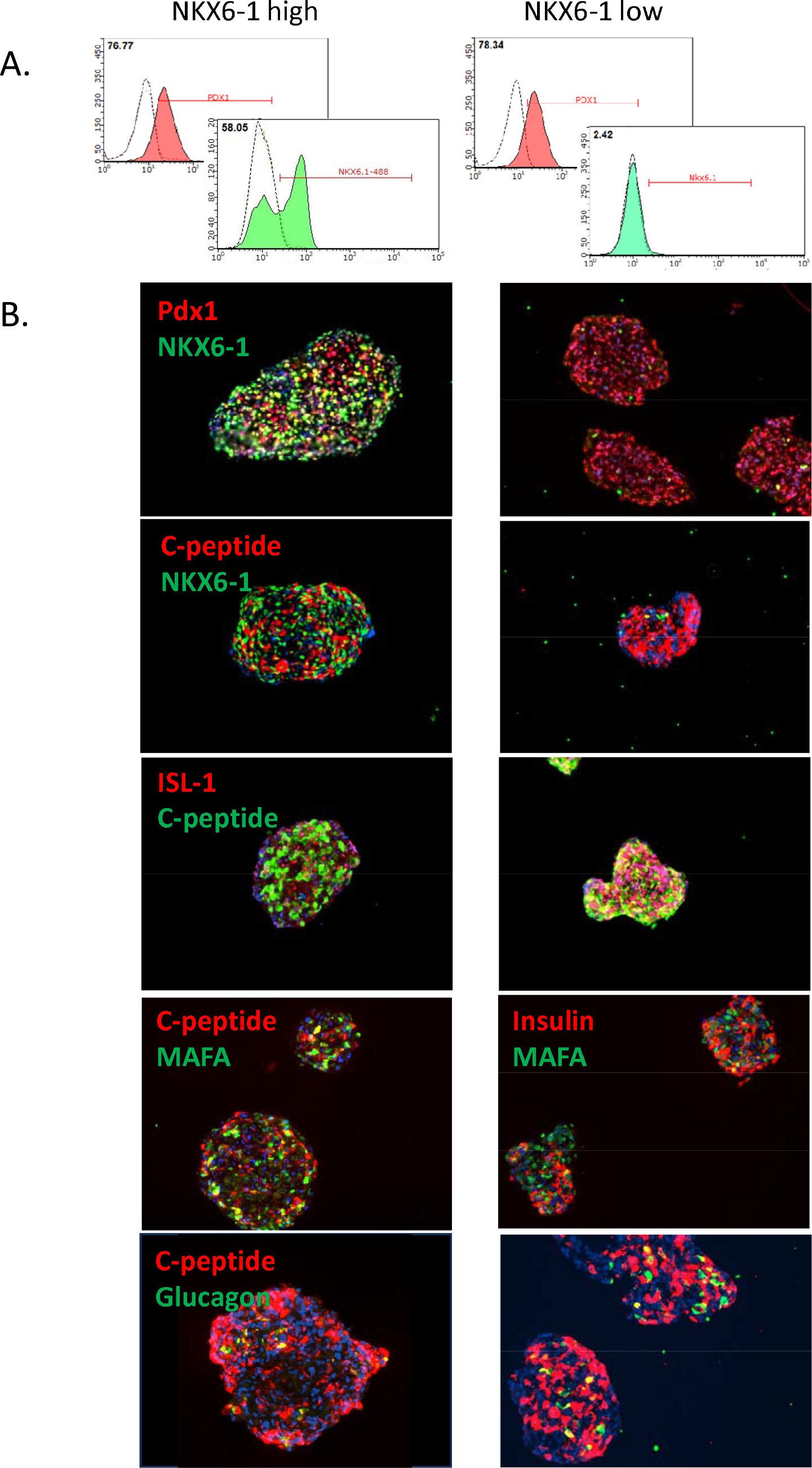
Flow cytometry (A) and immunofluorescent images (B) of NKX6-1 high and low clusters. Day 30-35 clusters from each protocol were characterized for expression of key pancreatic islet transcription factors (Pdx1, Nkx6.1, ISL-1, MAFA) and hormones (C-peptide, glucagon). Clusters were similar except for expression of Nkx6.1.

**Figure 3.**
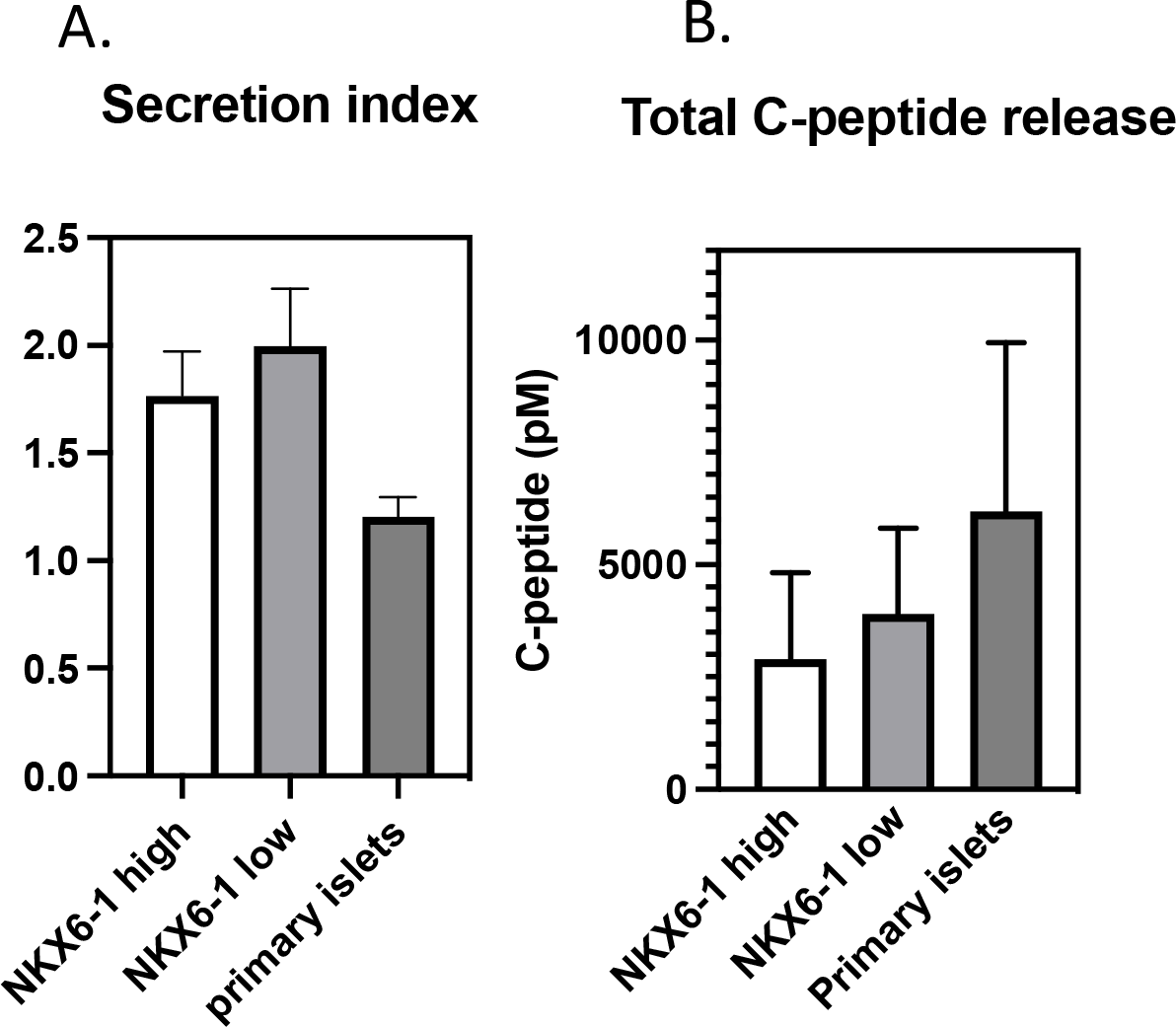
The secretion index (C-peptide secreted during static exposure to 16.7mM glucose/C-peptide secreted during exposure to 2.2mM glucose) of islet-like clusters is above 1 for islet like clusters derived from NKX6-1 low cultures (26 batches), NKX6-1 high cultures (16 batches) and primary human islets (6 preparations), indicating glucose sensitivity. Total C-peptide release that includes exposure to KCl was consistently higher in primary human islets, though not statistically significant.

Second differentiation protocol: We designed a second protocol that generated high proportions of cells that expressed PDX-1 and C-peptide without robust expression of NKX6-1. Flow cytometry and immunofluorescent staining (figure 2) confirmed a population of cells that co-express PDX-1 and insulin, and a low proportion of cells that express NKX6-1. This population more robustly expresses the transcription factors ISL1 and MAFA. ISL1 is expressed in early pancreatic endocrine progenitors and remains expressed in the mature pancreatic endocrine cells [27, 28]. ISL1 is a key activator of NKX6-1 and other genes found in mature glucose-sensing beta cells [9]. MAFA is a regulator of beta cell maturation [9]. The clusters from this protocol secreted C-peptide at levels similar to native islets in static culture conditions (Figure 3).

### In vivo performance

Islet like clusters from NKX6-1-high and NKX6-1-low populations were transplanted to immune-compromised mice and rats chemically rendered diabetic by application of streptozotocin. Clusters were delivered to the renal subcapsular space, intrasplenic, to a pouch formed from the omentum (rats), or a pouch formed from the gonadal fat pad (mice). Animals that received NKX6-1-low clusters reproducibly achieved glucose control within the time frames analyzed (figure 4A), while animals that received NKX6-1-high clusters did not (Figure S2). Glucose control was not influenced by site of implant at these sites. Glucose control was achieved in mice that received doses of NKX6-1-low clusters equivalent to 2,500 and 5,000 human islets. Animals demonstrating glucose control at 11-13 weeks post-engraftment demonstrated the ability to control blood glucose in a manner similar to non-diabetic controls after oral gavage of dextrose (figure 4B).

**Figure 4.**
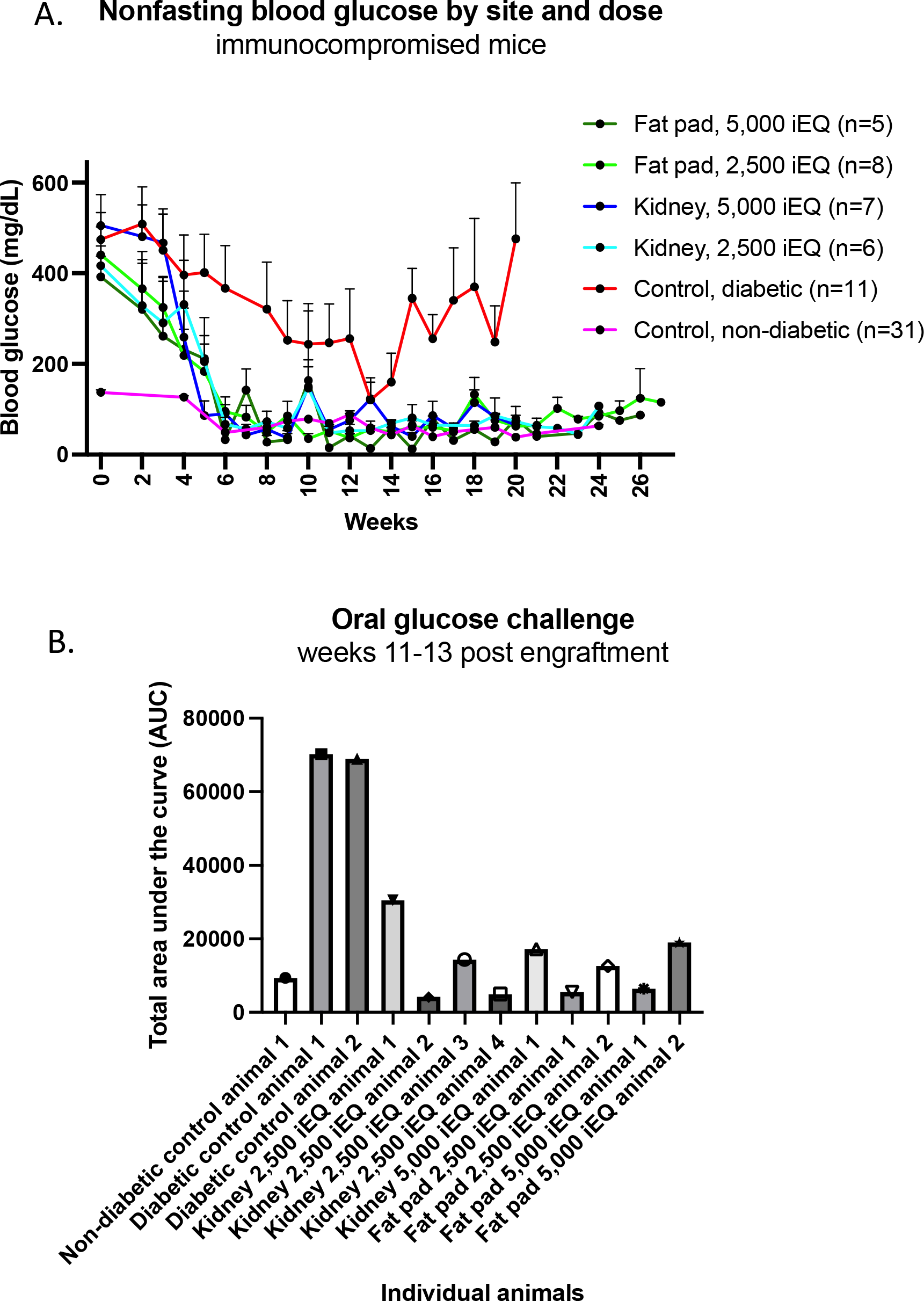
(A) Blood glucose of immune-compromised, STZ-induced diabetic mice was restored to levels similar to non-diabetic controls 5 weeks after implant of 2,500 (low) or 5,000 (high) islet equivalents to the renal subcapsular space or the fat pad. (B) Total area under the curve measurements of blood glucose following oral glucose challenge shows that treated animals, but not diabetic control animals, are capable of lowering blood glucose similarly to non-diabetic controls.

Grafts of NKX6-1-high clusters (first protocol) revealed a loss of C-peptide expression in the NKX6-1 positive population over time (figure 5A-D). This contrasts with previous reports that PDX-1/NKX6-1 co-positive cells mature in a hyperglycemic background in vivo to generate insulin-secreting cells homologous to native beta cells in this time frame [3, 16] Clusters from both protocols were found on explant to have been remodeled with host tissue to form structures with abundant, human endocrine tissue comingled with remodeled host tissue (figure 5 A-H). Although the clusters were delivered as a bolus, explants revealed islet like structures that were separated from each other by tissue that resembled ducts and parenchyma. Among explants of NKX6-1-low grafts, expression of endocrine markers within the islet-like structures were similar to native islets with robust chromogranin A (E) and PDX-1 (G) co-expressed with C-peptide or insulin. NKX6-1 was expressed with ISL1 and C-peptide, indicating in vivo activation. Pancreatic endocrine cells were also recognized by anti-human mitochondria antibody, demonstrating human origin. These data demonstrate that pancreatic clusters derived from SR1423 stem cells actively cooperate or drive remodeling of rodent host tissue to form stable pancreatic grafts with anatomic features of endocrine pancreas. Clusters from NKX6-1-low population maintain persistent expression of c-peptide, continue along islet maturation pathways and are potently capable of regulating rodent blood glucose.

**Figure 5.**
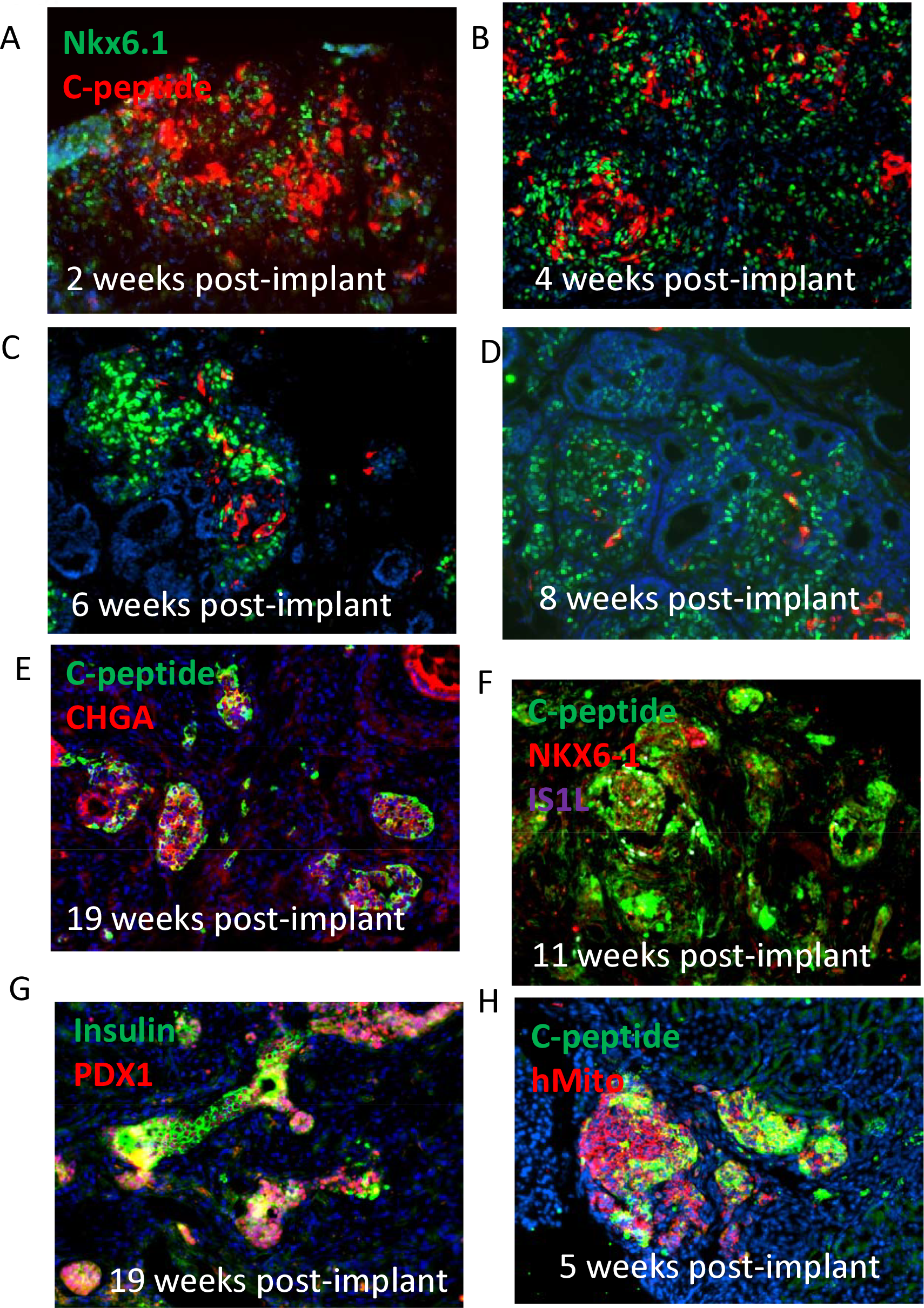
Tissue section immunofluorescent staining of explanted grafts at various time points demonstrate a remodeled tissue with elements of functional endocrine pancreas. NKX6-1 expressing cells from NKX6-1-high grafts demonstrated gradual loss of C-peptide expression over the first 8 weeks post implant (A-D). NKX6-1 low grafts maintained persistent expression of C-peptide or insulin (E,G), CHGA (E), PDX1 (G) and activated expression of NKX6-1 (F). Pancreatic endocrine cells co-expressed human mitochondria antigen (hMito) (H).

### NKX6-1 low characterization

Unsupervised transcriptome analysis of the NKX6-1-low protocol by scRNA seq using the Uniform Manifold Approximation and Projection (UMAP) method for dimensionality reduction reveals that stages 0 (undifferentiated), 1 and 2 cluster separately in discrete groupings in agreement with the drastic morphological changes that occur when stem cells launch differentiation (figure 6). These and subsequent stages manifest gene expression profiles consistent with progressive maturation to the pancreatic endocrine identity. Stages 1 and 2 exhibit endodermal gene expression profiles including CXCR4, SOX17, CER1, and LEFTY1 [29]. The guidance cues of stages 3-6 drive expression of genes characteristic of pancreatic commitment such as FOXA2, HNF4A, HNF1B, PDX-1, SOX9, AND PTF1A. The guidance cues of stages 7, 8, and 9 promote the progressive adoption of gene expression characteristic of the endocrine pancreas including, NKX2-2, NEUROG3, NEUROD1, MAFB, MAFA, PAX4, PAX6, ISL1, CHGA, and CHGB. The stage 9 cluster is adjacent to the cluster formed by native human islets. These findings of differentiation towards pancreatic islets are additionally supported by pseudotemporal trajectory analysis performed using monocle [30] which demonstrates a progressive, off-target-free differentiation of SR1423 stem cells into islet-like clusters. Furthermore, pathway analysis using VISION [31] demonstrates a stage-dependent acquisition of β cell-like identity and regulation of gene expression. Together, these data describe a progressive differentiation from stem cells through endoderm, gut tube, pancreatic progenitor and finally pancreatic endocrine commitment.

**Figure 6.**
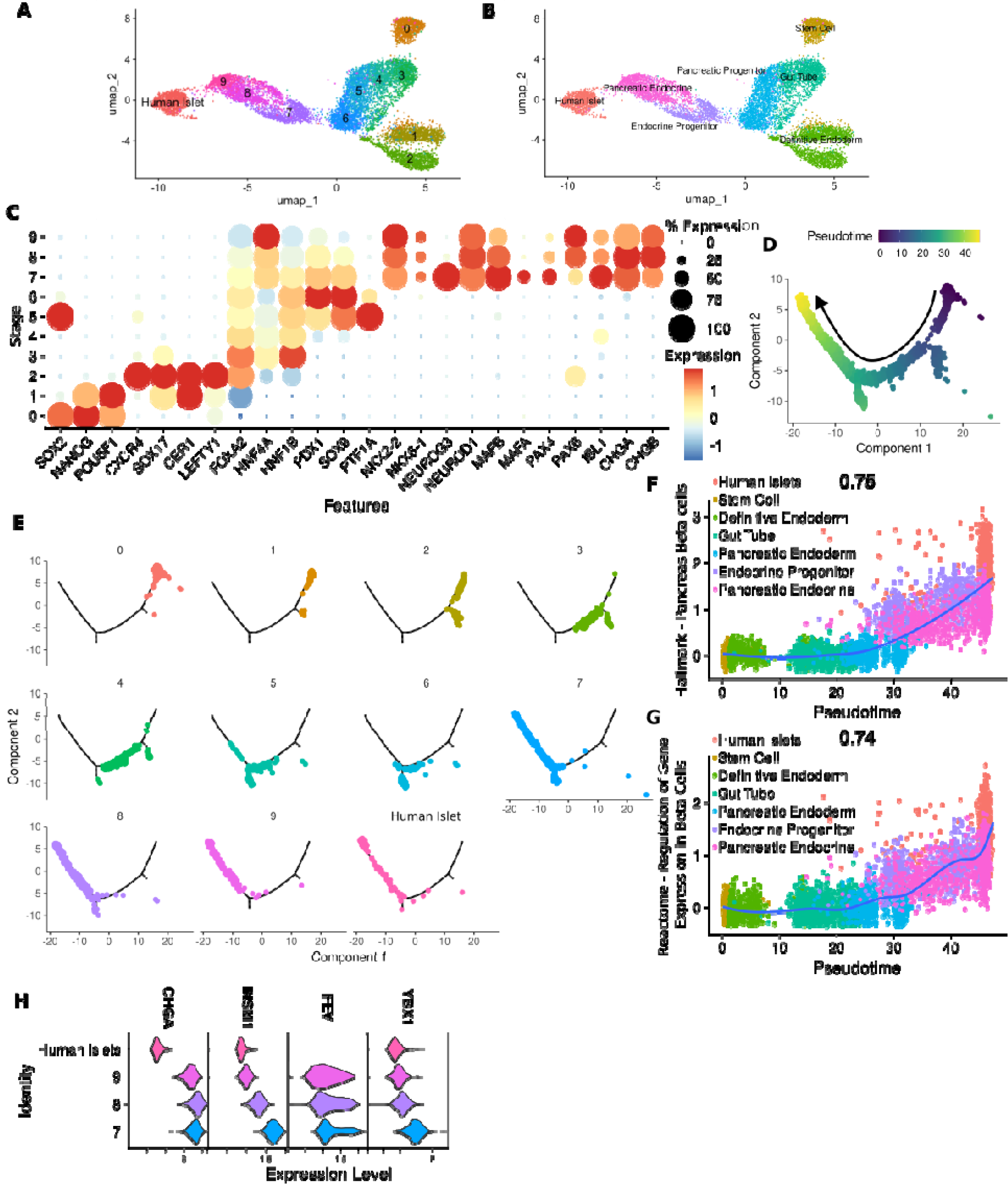
Transcriptome analysis of NKX6-1-low clusters and differentiation stages. (A) UMAP embedding of SR1423 stem cells, stages 1-9 of differentiation and human islets. Dimensionality reduction based on global reactome pathway scoring determined by VISION. (B) Grouping of SR1423 differentiation stages by pancreatic differentiation phases. (C) Dot plot of key genes associated with pancreatic differentiation phases from B. (D) Differentiation trajectory and pseudotime determine by monocle. (E) Differentiation trajectory split by differentiation stage. Projection of Pancreas Beta cell (F) and Regulation of Gene Expression in Beta Cells gene signatures in pseudotime. (H) Expression of CHGA, INSM1, FEV, and YBX1 in stage 7-9 and human islet cells.

Next, we sought to further characterize the late stages of SR1423 differentiation by focusing our analyses on pancreatic endocrine cells. Based on our snRNA-seq data, SR1423 differentiation yields a pure population of endocrine cells exhibiting 100% positivity for CHGA among pancreatic endocrine and endocrine progenitors (stages 7-9, figure 6H). YBX1-INSM1-FEV is required for the specification and maintenance of endocrine progenitor identity [9]. These genes are also found in stages 7-9 as well as mature human islets. Despite glucose responsive insulin production these cells retain expression of genes critical to maintenance of pancreatic progenitors affirming SR1423-derived pancreatic endocrine cells as less mature than adult islets but committed to that fate.

### Conclusion

In conclusion, these data suggest that multiple stem-cell derived pancreatic populations can be produced, and that these populations have disparate potency at controlling blood glucose in vivo. Furthermore, the profile of the stem cell derived pancreatic population that most closely aligns with developing states of the immature human pancreas is not fully defined. Seraxis pancreatic clusters are a highly pure population of pancreas-committed cells that are capable of potently controlling blood glucose in mouse and rat models of insulin-dependent diabetes within weeks of implant. We conclude that these clusters have therapeutic potential for physiologic control of blood glucose for insulin-requiring diabetes. A culture process that incorporates GMP compliant practices was designed to manufacture pancreatic clusters from SR1423 stem cells. Characterization and lot release criteria encompassing appropriate elements of consistency, purity and safety were added to comply with acceptable clinical practice.

## Discussion

In vitro, a widely accepted measure of endocrine potency is glucose stimulated insulin secretion. This report demonstrates that two disparate populations of cells with pancreatic identity, distinguished by expression of NKX6-1, have similar in vitro potency but different in vivo potency. Several protocols for causing the differentiation of pluripotent stem cells to pancreatic cells that express NKX6-1 are described in the scientific literature [2]. The population of NKX6-1 expressing cells described here are not claimed to be identical to NKX6-1 expressing cells derived from pluripotent cells and described by other groups [4, 14, 26, 32, 33]. It is unknown whether the monohormonal insulin-expressing cells that lost C-peptide expression in vivo de-differentiated, trans-differentiated to another endocrine phenotype, or would eventually have completed maturation and restarted insulin expression. In this context, however, a presumed in vitro marker of in vivo potency, expression of NKX6-1, was not reliable.

Animal implants of stem cell derived islets have been shown to take weeks longer to regulate blood glucose than native islets, despite showing glucose sensitive insulin secretion in vitro. This delay implies that stem cell derived islets are not functionally mature in vitro but can reach functional maturation in vivo. The hypothesis that only a small proportion of the progenitors are capable of in vivo maturation and that underdosing explains the delay to glucose regulation has been proposed [34]. Transcriptome profiles shown here demonstrates that stem cell derived pancreatic clusters share most homology with prenatal pancreatic endocrine populations [9]. Insulin expression and secretion in stem cell derived populations is therefore promiscuous and may not be a reliable measure of beta cell commitment [9]. Others have reached a similar conclusion in the context of cells co-expressing insulin and other pancreatic hormones [34, 35].

NKX6-1-low populations were able to consistently regulate blood glucose after 6 weeks. Significant tissue remodeling occurs within the first few weeks of each implant site evaluated (subcapsular renal space, intra splenic, and intra-abdominal omentum or fat pad). In each of these sites, islet-like structures formed as well as duct-like structures that were not observed in the in vitro clusters. Tissue injury repair and regeneration processes, including survival response to hypoxia and cell stress, likely promote the formation of these structures from the implanted material [36]. It has been reported that in regenerating pancreata show improved islet growth and higher insulin content near ducts [37]. Additionally, duct cells secrete angiogenic cytokines that would facilitate vascularization of the implanted site [38]. Similarly, others have demonstrated the benefits of promoting vascularization in poorly vascularized sites [39, 40], and of promoting pancreatic niche by delivering clusters in a synthetic or natural scaffold [41, 42]. Tissue remodeling may therefore be key in the establishment of a functional niche in which beta cell maturation occurs. Our interpretation of explants is that the remodeling process is a critical determinant of graft function and therefore impacts the effective dose.

Transplantation of islet-like clusters to insulin-requiring diabetics in a clinical setting could be improved by enhancing the allogeneic compatibility of the graft. That goal can be achieved by disrupting expression of major histocompatibility proteins I and II, the main antigenic drivers of allogeneic tissue rejection [43]. Our laboratory has therefore performed these and other edits on SR1423u defining new cell lines to improve tolerability and safety for future use in non-immunosuppressed recipients.

The islet like clusters described here are the subject of characterization and non-clinical studies to support regulatory filing for clinical testing. The cell expansion and differentiation protocol have been adapted to a manufacturing process following cGMP guidelines using closed, scalable vessels within a cleanroom environment. This work also demonstrates progress with the potential to achieve a potent and safe islet replacement therapy for insulin requiring diabetes.

## Methods

### hPSC Scorecard

Undifferentiated SR1423u cells from the Seraxis working cell bank were thawed and routinely cultured until passage 23. Near-confluent 10 cm dishes were lysed in TriZOL and stored at -80C. RNA was harvested from the TriZOL lysate according to the purification protocol described in the ThermoFisher TaqMan hPSC Scorecard manual. The nucleic acid pellets were dissolved in 50uL of nuclease-free water, then aliquots were stored at -80C. Once thawed, nucleic acid concentration was determined by NanoDrop. Reverse transcription was performed in duplicate following the Thermo Fisher TaqMan hPSC Scorecard Assay protocol. Briefly, 2ug of nucleic acid was mixed with ThermoFisher High Capacity cDNA synthesis reagents (random hexamer primers, RNase inhibitor, buffer, reverse transcriptase, dNTPs, water). The mix was incubated at 37C for 2 hours, then enzymes were heat-inactivated at 85C for 5 minutes. cDNA was stored at - 80C until use. Quantitative PCR was performed according to the ThermoFisher hPSC Scorecard assay manual on 10/25/2021. Pre-made ThermoFisher hPSC Scorecard panel 96 well plates were loaded with cDNA template and TaqMan Fast Advanced PCR Master Mix. Cycling parameters were programmed according to the Scorecard assay manual. The procedure was repeated. The Excel results files contained Ct values for each well. These values were then copied into an Excel template provided by ThermoFisher which has been formatted for use with the Scorecard online software. The formatted values were saved as .txt files for uploading into the Scorecard online software. Results from each replicate were uploaded into the software and pooled.

### Cell culture and Differentiation

Undifferentiated stem cells were maintained in mTeSR-E8 (Stem Cell Technologies) on plastic dishes precoated with full-length vitronectin (Peprotech) with daily media changes.

Guidance cues for differentiation and their durations are outlined in table 1. Stages 1 and 2 used basal media comprised of 1:1 mixture RPMI and DMEM F-12 supplemented with BSA and SM1 (Stem Cell Technology) or B27 (ThermoFisher) Stages 3-7 used basal media of DMEM supplemented with BSA and SM1 (Stem Cell Technology) or B27 (ThermoFisher). Stages 7-9 used a 1:3 mixture of CMRL and RPMI basal media supplemented with BSA. Stages 6-9 were carried out in suspension culture using a vertical wheel bioreactor culture system.

### Flow Cytometry

Single cell suspensions were prepared from clusters incubation in trypsin/EDTA (VWR). 20% of the total volume of fetal bovine serum was added and the tube placed on ice. The suspension was passed through a 70um filter. The eluate was quantified from 20ul aliquots that were mixed with trypan blue and counted twice using the TC20 automated cell counter (BioRad). Cells were pelleted and resuspended in 1 mL of Cytofix/Cytoperm solution (Becton Dickinson) and incubated at ambient temperature for 30 minutes. 150,000 cells were pelleted in a refrigerated centrifuge and wash in BD wash buffer (Becton Dickinson). Cells were pelleted and resuspended in PhosPerm Buffer III (Becton Dickinson) for 20-30 minutes on ice. Cells were pelleted and resuspended in 1:100 dilution of fluorophore conjugated secondary antibodies in BD wash buffer and incubated for 1 hour at room temperature or overnight at 4C. Cells were rinsed in BD wash buffer and resuspended in 1%BSA in PBS. Samples were quantified alongside unstained controls using a Guava flow cytometry system.

### Immunofluorescence

Samples were fixed for 30 minutes in 4% paraformaldehyde, rinsed in PBS and resuspended in 30% sucrose until sedimentation to the bottom of the vessel. Fixed samples were cryopreserved in OCT media cryosectioned to a thickness of 10um and collected on silane-coated microscope slides (VWR). Sections were permeabilized in 2.5% TX-100 (Sigma) dissolved in PBS for 20 minutes, rinsed and incubated in a blocking solution of 1% TX-100 and 10% horse serum for 1 hour. Sections were rinsed and incubated overnight at 4C in primary antibody diluted 1:100 in PBS supplemented with 1% TX-100 and 2% horse serum. Slides were rinsed and incubated in 1:1000 dilution of fluorophore conjugated secondary antibodies raised in horse for 1 hour. Sections were rinsed, incubated in 20nM Hoescht dye for 2 minutes, rinsed, secured under a coverslip and visualized using an Evos Fluorescent microscope (ThermoFischer).

### scRNA-seq

SR1423 differentiated cells were harvested on the final day of each differentiation stage. For adherent culture stages, one confluent 15cm tissue culture dish was washed with PBS then treated with 5mM EDTA for 5 minutes at 37C and single cell suspensions were generated by manual trituration. Single cells were cryopreserved prior to library preparation. For suspensions phases of SR1423 differentiation approximately 50mg of clusters were collected and cryopreserved in culture media + 10% DMSO. Nuclei were isolated directly from cryopreserved cell and clusters samples which were then used for single cell library preparation by Sigulomics, Inc using 10x Genomics 3’ library prep kit v3 chemistry. Nuclei were loaded into the chromium controller to target 5,000 nuclei for sequencing. Following library preparation, sequencing was performed at Novogene USA using the NovaSeq 6000 sequencer (Illumina) with a target of 35,000 reads per cell.

### scRNA-seq data processing and analysis

Following sequencing, reads were aligned to the human reference genome (GRCh38) using CellRanger v7.1.0 (10x Genomics). Next, data were processed in R V4.3.0 using Seurat v5. Filtering was performed to exclude cells with fewer than 500 genes expressed and less than 3000 transcripts detected. Following filtering data were normalized and the top 3000 variable genes were selected for integration. Principal component analysis was performed on these variable genes and data were integrated via reciprocal principal component analysis using the first 30 principal components. The RNA expression matrix was loaded into VISION v3.0.1 [31] and single cell pathway analysis was performed using default parameters and the full Reactome pathway database. The results of VISION pathway analysis were added to the Seurat object as a new assay. The top 200 variable pathways were identified and used for principal component analysis, nearest neighbor graph embedding, and clustering with a resolution of 0.5. Uniform manifold projection was calculated based on the first 20 principal components of the Reactome pathway analysis. Drop outs were imputed using the RunALRA function implemented through SeuratWrappers using default parameters. Drop out-imputed data were only used for visualizing gene expression.

Trajectory and pseudotime analyses were performed using Monocle v2 [30]. For these analyses, the normalized RNA expression matrix was downsampled to 1000 cells from each starting sample then loaded into monocle. Within monocle, dimensionality reduction was performed using the DDRTree method and pseudotime computed.

### Animal experiments

Animal husbandry and animal implants, care and monitoring were carried out using IACUC-approved protocols at the contract research facilities of Noble Life Sciences and Bioqual, Inc. NSG mice were sourced from Jackson Labs. SRG rats were sourced from Envigo.

#### Streptozotocin

Diabetes was induced with a single intraperitoneal injection of 150mg/kg of streptozotocin (mice) (Sigma) or 50mg/kg (rats) to nonfasting animals. Diabetes was confirmed by two blood glucose measurements above 350mg/dL. Hyperglycemic animals were provided a dose of subcutaneous insulin pellets following manufacturer’s instructions (Linshin Canada).

#### Implant

Subcapsular renal implant. 2,500 or 5,000 islet equivalents of Seraxis islet-like clusters were counted using the islet cell counter (Biorep) automated method and transferred to a 1.5ml conical tube for mouse implants. 10,000 or 40,000 clusters were manually counted and transferred to a 1.5ml conical tube for rat implants. Islet like clusters were pelleted using a tabletop microfuge and aspirated into the tip of a 15-gauge flexible gavage needle. This was deposited into a renal subcapsular pouch using standard veterinary surgical techniques.

Fat pad/omentum implants. The gonadal fat pad was exited from the lower abdomen using standard veterinary surgical technique and spread flat along the abdomen. 2,500 or 5,000 islet equivalents of Seraxis clusters were pelleted using a tabletop microfuge and resuspended in 20ul of purified human plasma (Tisseel, component A). The cluster/plasma mixture was deposited onto the fat pad. An equal volume of fibrin solution (Tisseel, component B) was layered on to the cluster plasma mixture, which was enclosed by folding of the remaining fat pad tissue. For rat implants, the omentum was exited from the upper abdomen and 10,000 or 40,000 clusters were deposited in the same fashion.

Intrasplenic implant. 10,000 or 40,000 islet like clusters were pelleted and aspirated into the bore of a 20-gauge hypodermic syringe. The rat spleen was exited through a dorsal abdominal incision. The syringe was inserted to the spleen longitudinally and slowly vacated as it was withdrawn.

### Glucose sensitive insulin secretion

50 islet-like clusters or primary islets were incubated sequentially in buffer supplemented with 2.2mM and 16.7mM mM glucose for 30minutes, followed by a 30 minute incubation in KCl. Supernatant was analyzed for C-peptide content by ELISA (Mercodia). The secretion index represents the ratio of C-peptide secreted during exposure to 16.7mM glucose over C-peptide secreted during exposure to 2.2mM glucose. Total C-peptide release corresponds to the sum of C-peptide secreted during both exposures and KCl.

## Acknowledgements

Snowden, A., Bloom, J., Hosseini, M., Rodman, L., McWilliams, I., Tran, C., and Williams, K. contributed to the manufacture, testing, and lot release of pancreatic clusters.

## Figure Legends

**Figure S1.**
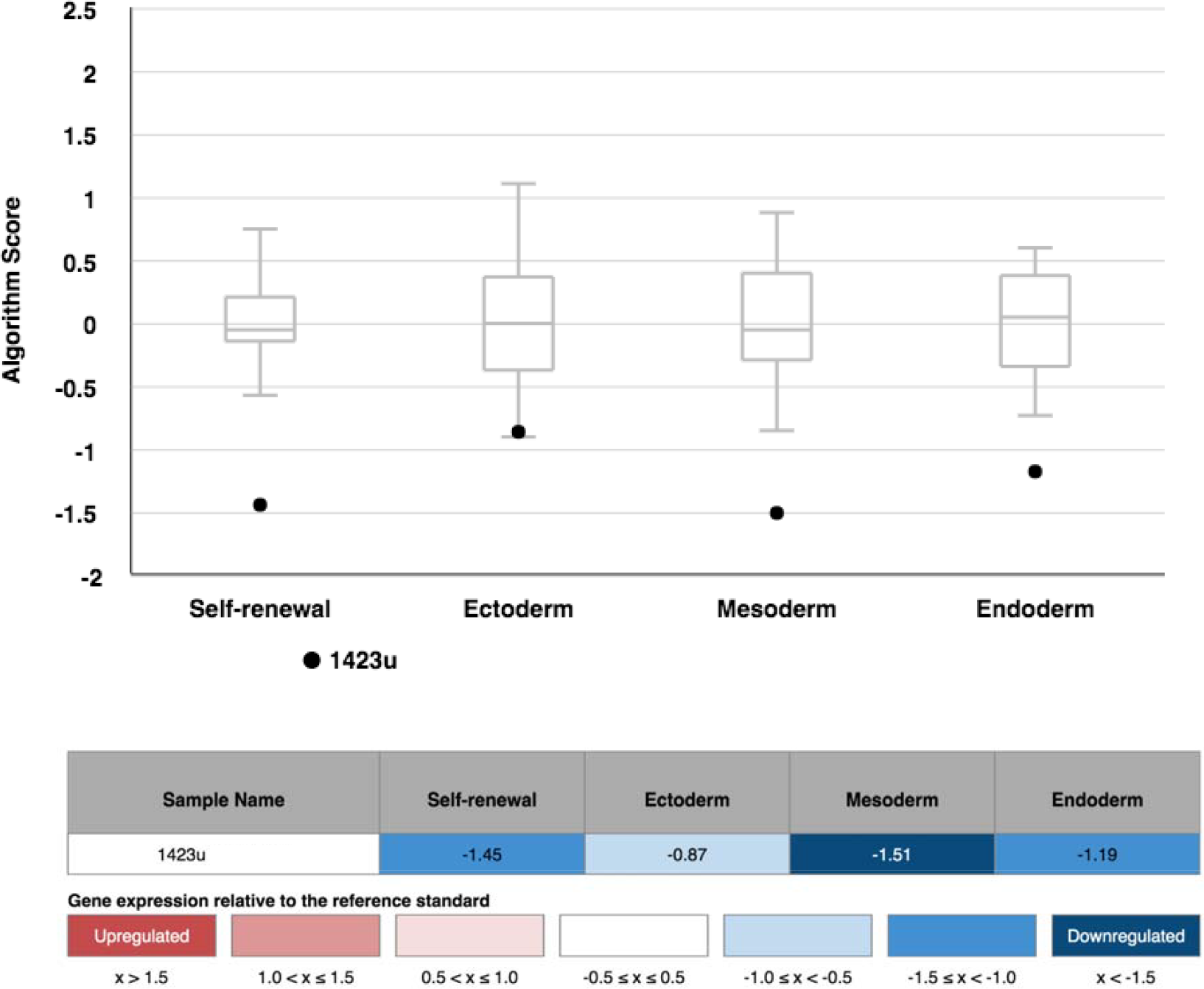
SR1423 RNA was analyzed by Scorecard qPCR for the expression of a panel of 94 genes associated with pluripotency, endoderm, mesoderm and ectoderm following the Scorecard instructions. The datasets were formatted for the Thermo Fisher TaqMan hPSC Scorecard Assay and uploaded to the hPSC Scorecard analysis software. Box and whisker plots represent reference gene datasets. Results reveal a transcriptome with a differentiation expression profile outside of the significant range of the mean of the reference dataset. Genes related to mesodermal differentiation were the most severely downregulated, correlating with inability for mesodermal differentiation in vitro.

**Figure S2.**
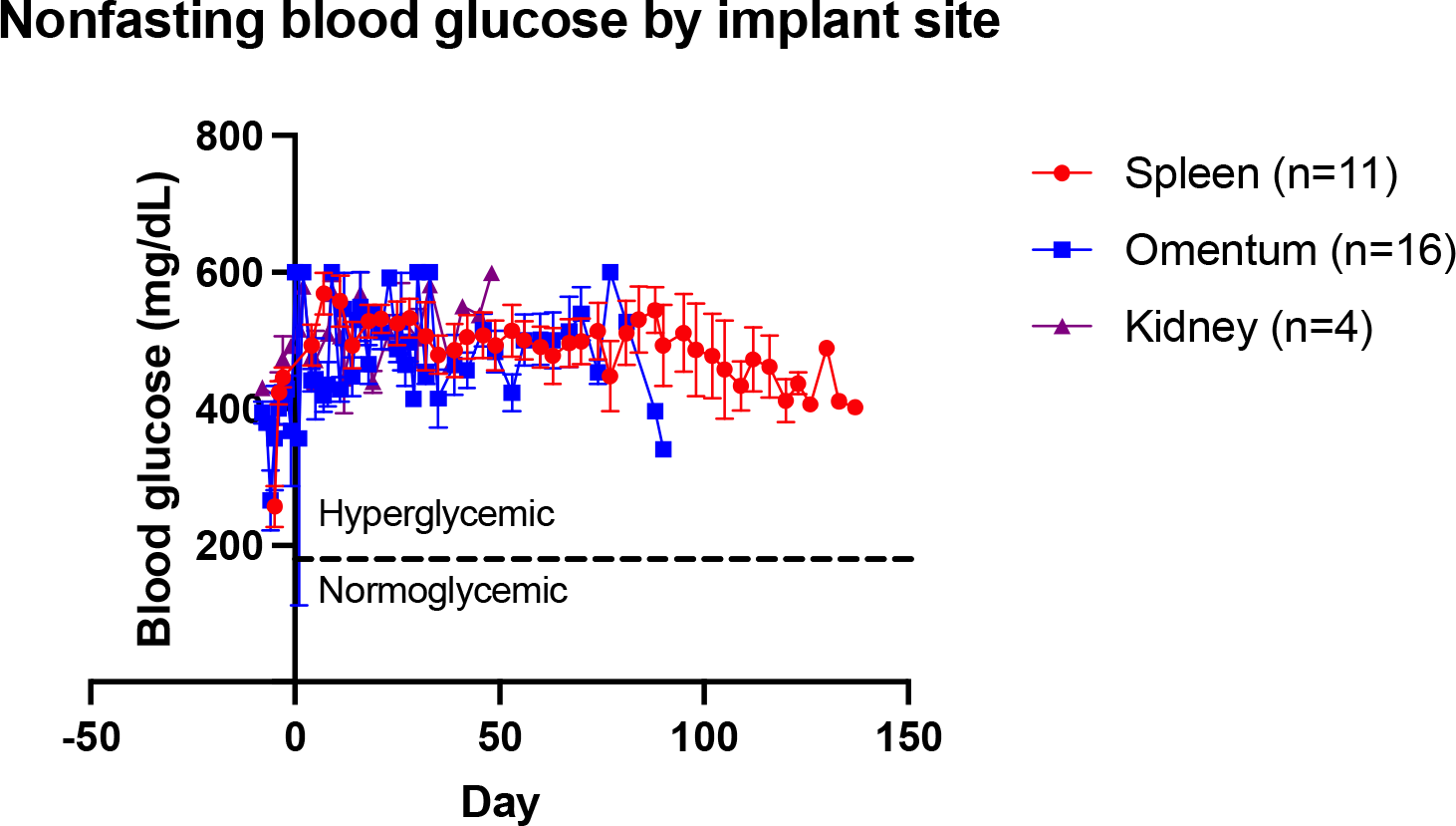
10,000-40,000 Nkx6.1 rich islet like clusters did not control blood glucose after implant to STZ induced, diabetic immune compromised rats. Implant site did not influence glucose control.

## References

1. Witkowski, P., et al., The demise of islet allotransplantation in the United States: A call for an urgent regulatory update. Am J Transplant, 2021. 21(4): p. 1365–1375.

2. Hogrebe, N.J., M. Ishahak, and J.R. Millman, Developments in stem cell-derived islet replacement therapy for treating type 1 diabetes. Cell Stem Cell, 2023. 30(5): p. 530–548.

3. Augsornworawat, P., et al., Single-nucleus multi-omics of human stem cell-derived islets identifies deficiencies in lineage specification. Nat Cell Biol, 2023. 25(6): p. 904–916.

4. Balboa, D., et al., Functional, metabolic and transcriptional maturation of human pancreatic islets derived from stem cells. Nat Biotechnol, 2022. 40(7): p. 1042–1055.

5. Veres, A., et al., Charting cellular identity during human in vitro beta-cell differentiation. Nature, 2019. 569(7756): p. 368–373.

6. International Stem Cell, I., Assessment of established techniques to determine developmental and malignant potential of human pluripotent stem cells. Nat Commun, 2018. 9(1): p. 1925.

7. Southard, S.M., R.P. Kotipatruni, and W.L. Rust, Generation and selection of pluripotent stem cells for robust differentiation to insulin-secreting cells capable of reversing diabetes in rodents. PLoS One, 2018. 13(9): p. e0203126.

8. Zhu, H., et al., Understanding cell fate acquisition in stem-cell-derived pancreatic islets using single-cell multiome-inferred regulomes. Dev Cell, 2023. 58(9): p. 727–743 e11.

9. Ma, Z., et al., Deciphering early human pancreas development at the single-cell level. Nat Commun, 2023. 14(1): p. 5354.

10. Ebrahim, N., K. Shakirova, and E. Dashinimaev, PDX1 is the cornerstone of pancreatic beta-cell functions and identity. Front Mol Biosci, 2022. 9: p. 1091757.

11. Aigha, II and E.M. Abdelalim, NKX6.1 transcription factor: a crucial regulator of pancreatic beta cell development, identity, and proliferation. Stem Cell Res Ther, 2020. 11(1): p. 459.

12. Memon, B. and E.M. Abdelalim, Stem Cell Therapy for Diabetes: Beta Cells versus Pancreatic Progenitors. Cells, 2020. 9(2).

13. Pellegrini, S., et al., Transcriptional dynamics of induced pluripotent stem cell differentiation into beta cells reveals full endodermal commitment and homology with human islets. Cytotherapy, 2021. 23(4): p. 311–319.

14. Velazco-Cruz, L., et al., Acquisition of Dynamic Function in Human Stem Cell-Derived beta Cells. Stem Cell Reports, 2019. 12(2): p. 351–365.

15. Egozi, A., et al., Insulin is expressed by enteroendocrine cells during human fetal development. Nat Med, 2021. 27(12): p. 2104–2107.

16. Augsornworawat, P., et al., Single-Cell Transcriptome Profiling Reveals beta Cell Maturation in Stem Cell-Derived Islets after Transplantation. Cell Rep, 2020. 32(8): p. 108067.

17. Du, W., et al., Pharmacological conversion of gut epithelial cells into insulin-producing cells lowers glycemia in diabetic animals. J Clin Invest, 2022. 132(24).

18. Melton, D., The promise of stem cell-derived islet replacement therapy. Diabetologia, 2021. 64(5): p. 1030–1036.

19. Singh, R., et al., Enhanced structure and function of human pluripotent stem cell-derived beta-cells cultured on extracellular matrix. Stem Cells Transl Med, 2021. 10(3): p. 492–505.

20. Hansson, M., et al., Artifactual insulin release from differentiated embryonic stem cells. Diabetes, 2004. 53(10): p. 2603–9.

21. Jones, A.G. and A.T. Hattersley, The clinical utility of C-peptide measurement in the care of patients with diabetes. Diabet Med, 2013. 30(7): p. 803–17.

22. Bock, C., et al., Reference Maps of human ES and iPS cell variation enable highthroughput characterization of pluripotent cell lines. Cell, 2011. 144(3): p. 439–52.

23. Galdos, F.X., et al., Combined lineage tracing and scRNA-seq reveals unexpected first heart field predominance of human iPSC differentiation. Elife, 2023. 12.

24. Weng, C., et al., Single-cell lineage analysis reveals extensive multimodal transcriptional control during directed beta-cell differentiation. Nat Metab, 2020. 2(12): p. 1443–1458.

25. Wang, L., et al., NANOG and LIN28 dramatically improve human cell reprogramming by modulating LIN41 and canonical WNT activities. Biol Open, 2019. 8(12).

26. Nostro, M.C., et al., Efficient generation of NKX6-1+ pancreatic progenitors from multiple human pluripotent stem cell lines. Stem Cell Reports, 2015. 4(4): p. 591–604.

27. Ediger, B.N., et al., Islet-1 Is essential for pancreatic beta-cell function. Diabetes, 2014. 63(12): p. 4206–17.

28. Sasaki, S., et al., Spatial and transcriptional heterogeneity of pancreatic beta cell neogenesis revealed by a time-resolved reporter system. Diabetologia, 2022. 65(5): p. 811–828.

29. Chu, L.F., et al., Single-cell RNA-seq reveals novel regulators of human embryonic stem cell differentiation to definitive endoderm. Genome Biol, 2016. 17(1): p. 173.

30. Qiu, X., et al., Reversed graph embedding resolves complex single-cell trajectories. Nat Methods, 2017. 14(10): p. 979–982.

31. DeTomaso, D., et al., Functional interpretation of single cell similarity maps. Nat Commun, 2019. 10(1): p. 4376.

32. Maxwell, K.G., et al., Differential Function and Maturation of Human Stem Cell-Derived Islets After Transplantation. Stem Cells Transl Med, 2022. 11(3): p. 322–331.

33. Pagliuca, F.W., et al., Generation of functional human pancreatic beta cells in vitro. Cell, 2014. 159(2): p. 428–39.

34. Pagliuca, F.W. and D.A. Melton, How to make a functional beta-cell. Development, 2013. 140(12): p. 2472–83.

35. Rezania, A., et al., Maturation of human embryonic stem cell-derived pancreatic progenitors into functional islets capable of treating pre-existing diabetes in mice. Diabetes, 2012. 61(8): p. 2016–29.

36. Fuchs, E. and H.M. Blau, Tissue Stem Cells: Architects of Their Niches. Cell Stem Cell, 2020. 27(4): p. 532–556.

37. Overi, D., et al., Islet Regeneration and Pancreatic Duct Glands in Human and Experimental Diabetes. Front Cell Dev Biol, 2022. 10: p. 814165.

38. Movahedi, B., et al., Pancreatic duct cells in human islet cell preparations are a source of angiogenic cytokines interleukin-8 and vascular endothelial growth factor. Diabetes, 2008. 57(8): p. 2128–36.

39. Vlahos, A.E., et al., Endothelialized collagen based pseudo-islets enables tuneable subcutaneous diabetes therapy. Biomaterials, 2020. 232: p. 119710.

40. Dang, H.P., et al., Cell delivery systems: Toward the next generation of cell therapies for type 1 diabetes. J Cell Mol Med, 2022. 26(18): p. 4756–4767.

41. Rojas-Canales, D., et al., Intracutaneous Transplantation of Islets Within a Biodegradable Temporizing Matrix as an Alternative Site for Islet Transplantation. Diabetes, 2023. 72(6): p. 758–768.

42. Coronel, M.M. and C.L. Stabler, Engineering a local microenvironment for pancreatic islet replacement. Curr Opin Biotechnol, 2013. 24(5): p. 900–8.

43. Montgomery, R.A., et al., HLA in transplantation. Nat Rev Nephrol, 2018. 14(9): p. 558–570.

